# mMap: a comprehensive and continuously updated species distribution database facilitating the understanding of mosquito diversity and distribution patterns

**DOI:** 10.64898/2025.12.22.695893

**Authors:** Xiang Guo, Shu Zeng, Haiyang Chen, Ziyao Li, Xiaohua Liu, Liu Ge, Qing He, Wenyi Li, Xiaohong Zhou

## Abstract

Mosquitoes, as vectors of deadly diseases like malaria and dengue, exhibit complex and dynamic global distribution patterns influenced by climate change and human activity. However, gaps in species-level data and biases in existing databases hinder accurate risk assessments. Here, we present mMap (v1.01), a comprehensive and curated mosquito distribution database integrating multi-source records from literature (36,242 entries across 1,809 studies), museum collections, and field reports in 6 languages. Systematic data harmonization and validation workflows address taxonomic and spatial uncertainties, enabling family-level analyses previously constrained by fragmented data. Our results reveal significant range expansions for 28 invasive species (e.g., *Aedes albopictus* in Europe) and corrections to historical distributions (e.g., *Ae. vexans* in 22 new countries). The interactive mMap platform (accessible at https://mos-map.vercel.app/) provides open-access tools for visualizing spatial-temporal trends, automated SDM forecasting, and expert-curated species profiles. By unifying dispersed data into a scalable framework, mMap advances research on mosquito biodiversity, invasion ecology, and disease risk mapping, offering a critical resource for global health strategies in an era of rapid environmental change.

## 1. Introduction

Mosquitoes are distributed globally, with the exception of Antarctica. Their remarkable reproductive capacity and adaptability to diverse habitats have contributed to their high species diversity[1]. The family Culicidae (Insecta, Diptera) is divided into two subfamilies, Culexinae and Anophelinae, encompassing 12 tribes, including Aedini and Anophelini, comprising a total of 63 genera and 3,147 species[2, 3]. Male mosquitoes typically feed on plant nectar, while females require blood meals for reproduction, often biting and causing irritation to humans and animals. As one of the most significant medical insects, mosquitoes serve as vectors for numerous severe mosquito-borne diseases, such as malaria, Zika virus disease, and dengue fever[4, 5]. It is estimated that approximately 700 million people worldwide are infected with various diseases transmitted by mosquitoes annually. These diseases exert a profound impact on public health, socioeconomic development, and ecological systems, posing a substantial threat to human survival. Furthermore, they jeopardize national security, social stability, and global safety, representing a critical challenge to the global community and all nations.

In recent years, urbanization, climate warming, and the intense fluctuations of the ENSO cycle have significantly altered the global ecological environment, thereby profoundly impacting the biodiversity and ecological behaviors of insects[6, 7, 8]. Notably, several mosquito species have exhibited marked global expansion trends[9, 10]. A prime example is *Aedes albopictus*, which has successfully adapted to urbanized environments and expanded its distribution range globally, spreading from its native regions in East Asia and Southeast Asia to all continents except Antarctica[10, 11]. Over the past two decades, *Ae. albopictus* has been recognized as one of the 100 most rapidly expanding invasive species worldwide[10]. Conversely, some mosquito species have experienced reductions in their habitat ranges. *Aedes aegypti* serves as a notable example; recent field ecological studies conducted across multiple regions in China indicate that this species has nearly disappeared in Guangdong and Hainan provinces, with only sporadic occurrences remaining in border areas of Yunnan adjacent to Myanmar [12, 13, 14]. Such discrepancies highlight the complexity of the ecological distribution and spatial structure of the Culicidae family, indicating that further research is needed to fully understand the overall biodiversity and global distribution patterns of mosquitoes.

Currently, well-studied mosquito species such as *Ae. albopictus* and *Anopheles sinensis* have established a relatively robust foundation for understanding their ecological distribution and spatial structure[11]. However, the biodiversity and global distribution patterns of many other mosquito species, as well as the Culicidae family as a whole, remain largely unreported. Given that it is impractical to have experts conduct in-depth research on every species, conducting comprehensive studies at the family level presents several challenges: (1) Data sources are fragmented, requiring the collection and integration of multi-source information from literature, museum collections, and specimen databases. Sole reliance on the single database (for example, GBIF) for ecological analysis may introduce significant bias; (2) Both data and metadata require thorough verification, updating, and analysis, which demands researchers possess a deep understanding of mosquito ecological habits and necessitates collaboration with expert panels for consultation and discussion. (3) At the family level, there is considerable variation in the quantity and quality of available data, compounded by the large number of species involved. To address this, an executable automated workflow must be developed to efficiently process and analyze the data. (4) Geographical disparities in field sampling intensity, driven by varying levels of research attention worldwide, must also be taken into account.

Previous ecological studies conducted at the family level in mammals and other invertebrates, such as ants[15, 16], have provided valuable technical references. In this study, we aim to integrate multi-source data to construct a highly diverse, comprehensive, and detailed distribution database for mosquito species. Additionally, we have developed a website, mMap, to visualize and present these findings. This research will not only provide robust evidence for understanding the diversity and global distribution patterns of mosquito species but also enhance our insights into the prevention and control of mosquito-borne infectious diseases.

## 2. Methods

### 2.1 Data Sources

We searched the databases relating to mosquito distribution in English, Chinese, Russ ian, Arabic, Japanese, and French up to 5th December, 2025 including PubMed (https://www.ncbi.nlm.nih.gov/pubmed/), the China National Knowledge Infrastructure Dat abases (CNKI, http://www.cnki.net), Wanfang (http://g.wanfangdata.com.cn/), NCL (ht tps://tpl.ncl.edu.tw/), J-STAGE (https://www.jstage.jst.go.jp/), Almanhal (https://almanhal.com/), Cyberleninka (https://cyberleninka.ru/) and Persee (https://www.persee.fr/). We implemented a strategy of performing separate searches by country/region, wherein different participants collaboratively conducted literature searches and data inclusion for the globally divided regions. For search terms, we used the following words: (“Country/area name” [Title]) AND ((Mosquito[Title]) OR (Aedes[Title]) OR (Culex [Title]) OR (Anopheles[Title])). All the original records of literature search were reco rded in Text S1.

The websites for WRBU (https://wrbu.si.edu/), Global Biodiversity Information Facility (GBIF) (https://www.gbif.org/), Global Mosquito Observations Dashboard (https://mosquitodashboard.org/) were also searched. Specialized books such as mosquito records from various regions in China and online reports from global governments were also reviewed.

### 2.2 Selection criteria

The selection criteria for the literature review were as follows: First, the authors were assigned to search and select literature, respectively, each step requiring double approval, and if any conflicts arose, they resolved them by consulting the senior authors. We also consulted some experts in the field when required. Second, after excluding duplicates, all literature was then screened by title, abstract and full-text (Text S1), followed by excluding the studies unrelated with mosquito distirbution. Third, those reports that were not specified by species identification of mosquito by authors were removed. But the relative records traced from the selected eligible reports’ references or the recommendation of the experts were added. All eligible studies concerning distribution of mosquito were then retained for the present descriptive analyses (Text S1).

### 2.3 Website architecture

The mMap website was implemented as a web application using TypeScript and React for front-end development. The core technologies include Next.js 14 (https://nextjs.org/) as the main framework with App Router architecture, @antv/l7 and LarkMap (https://l7.antv.antgroup.com/) for interactive geographic visualization, and Material React Table for data display. The user interface was built with TailwindCSS (https://tailwindcss.com/), NextUI (https://nextui.org/), and Material UI (https://mui.com/) component libraries. The application follows a serverless architecture with data stored in CSV format and served through Next.js API Routes, deployed on Vercel (https://vercel.com/) cloud platform. Extensive browser compatibility testing has been conducted on modern web browsers including Google Chrome, Firefox, Safari, and Microsoft Edge, ensuring a seamless user experience.

## 3. Results

### 3.1 Data source eligibility results

Our search of PubMed databases in English identified 24,653 potentially relevant records. Additional searches in Chinese databases, including CNKI and Wanfang, yielded a further 11,122 records. 226, 162, 78, and 1 records were were published in Russian, Spanish, Japanese, and Portuguese. Finally, 36,242 mosquito distribution records, comprising 27,632 original reports and 8,403 review-derived entries were included.

A comprehensive visualization of the mosquito distribution data compiled in the mMap database, highlighting its taxonomic, temporal, and geographic coverage was shown in Fig. 1. Taxonomically, *Aedes* is the most represented genus with 12,271 entries, followed by Anopheles (12,177, 33.60%) and Culex (8,809, 24.31%) (Fig. 1A). In contrast, rare genera such as Udaya and Zeugnomyia constitute only a negligible fraction of the dataset (Fig. 1A). Among all genera, Aedes, Anopheles, and Culex exhibit the highest species richness (Fig. 1B). The temporal trend of mosquito distribution records spans from the 1930s to the 2020s, showing exponential growth in records after 2000 (Fig. 1C). Africa (12,000 records) and Southeast Asia (10,000 records) are identified as the most data-rich regions, whereas country-level analysis highlights China, the United States, and Brazil as the nations with the highest documentation coverage (Fig. 1D).

**Figure 1.**
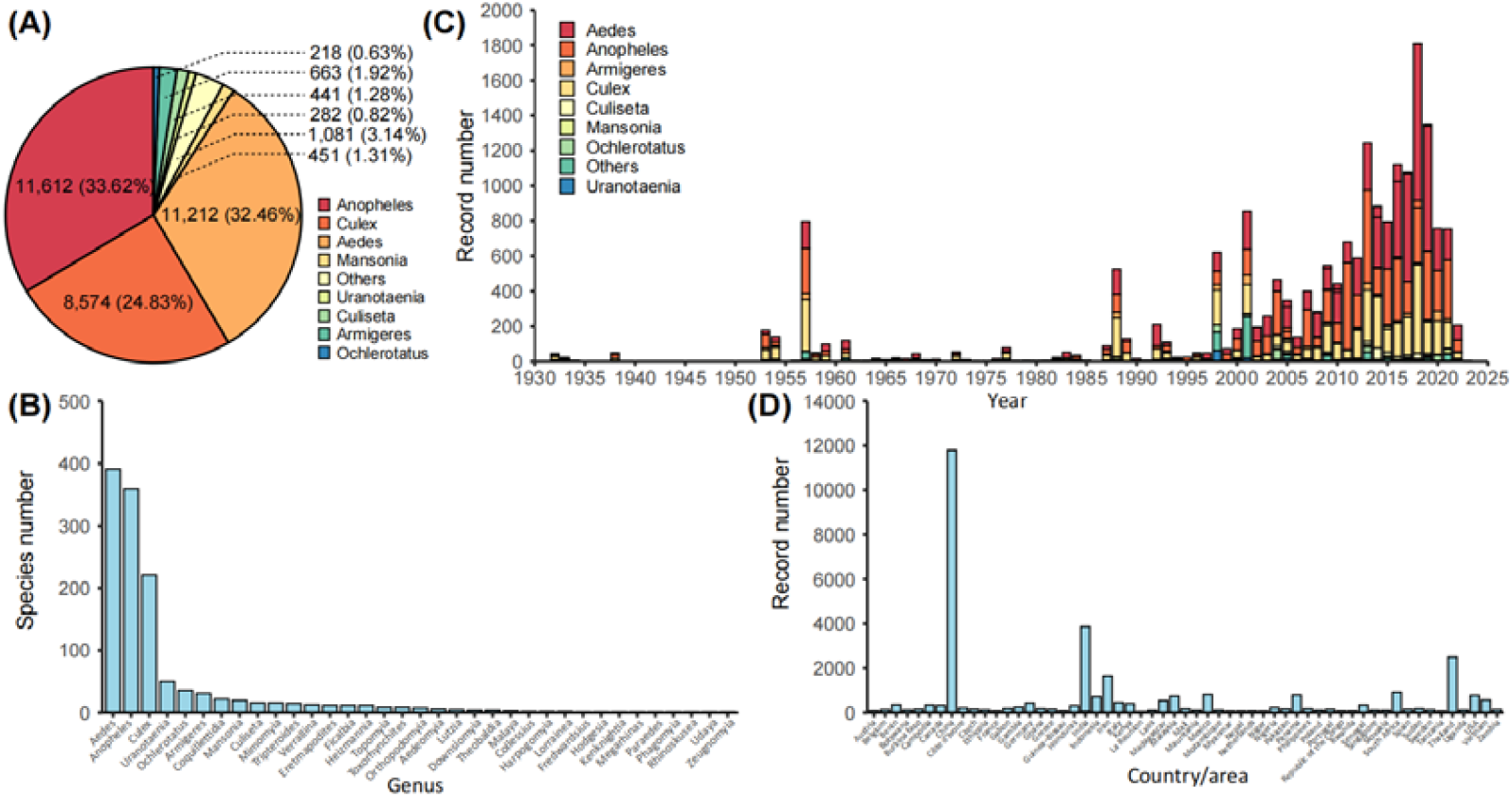
Taxonomic, temporal, and geographic coverage of the mMap database. (A) Genus-level composition of mosquito records, with Aedes, Anopheles, and Culex dominating the dataset. (B) Species richness across major genera, highlighting Aedes, Anopheles, and Culex as the most diverse. (C) Temporal trends in distribution records, showing exponential growth post-2000. (D) Geographic distribution of records at continental and country levels, with Africa and Southeast Asia as the most data-rich regions, and China, the United States, and Brazil as top-represented nations. (E) Leading journals contributing to mosquito distribution literature.

### 3.2 The mMap data supplements the global distribution knowledge of mosquito

The mMap data significantly enhances our understanding of the global distribution of mosquitoes (Fig. 2). Taking Aedes vexans as an example, mMap incorporates newly documented occurrences from 22 additional countries/regions, including previously unreported areas in West Africa, Southern Africa, Northeast Asia and Europe (Fig. 2A). Similar patterns are observed in other species such as *Ae. japonicus*, where mMap’s integrated records provide a more complete representation of their actual distributions (Fig. 2B).

**Figure 2.**
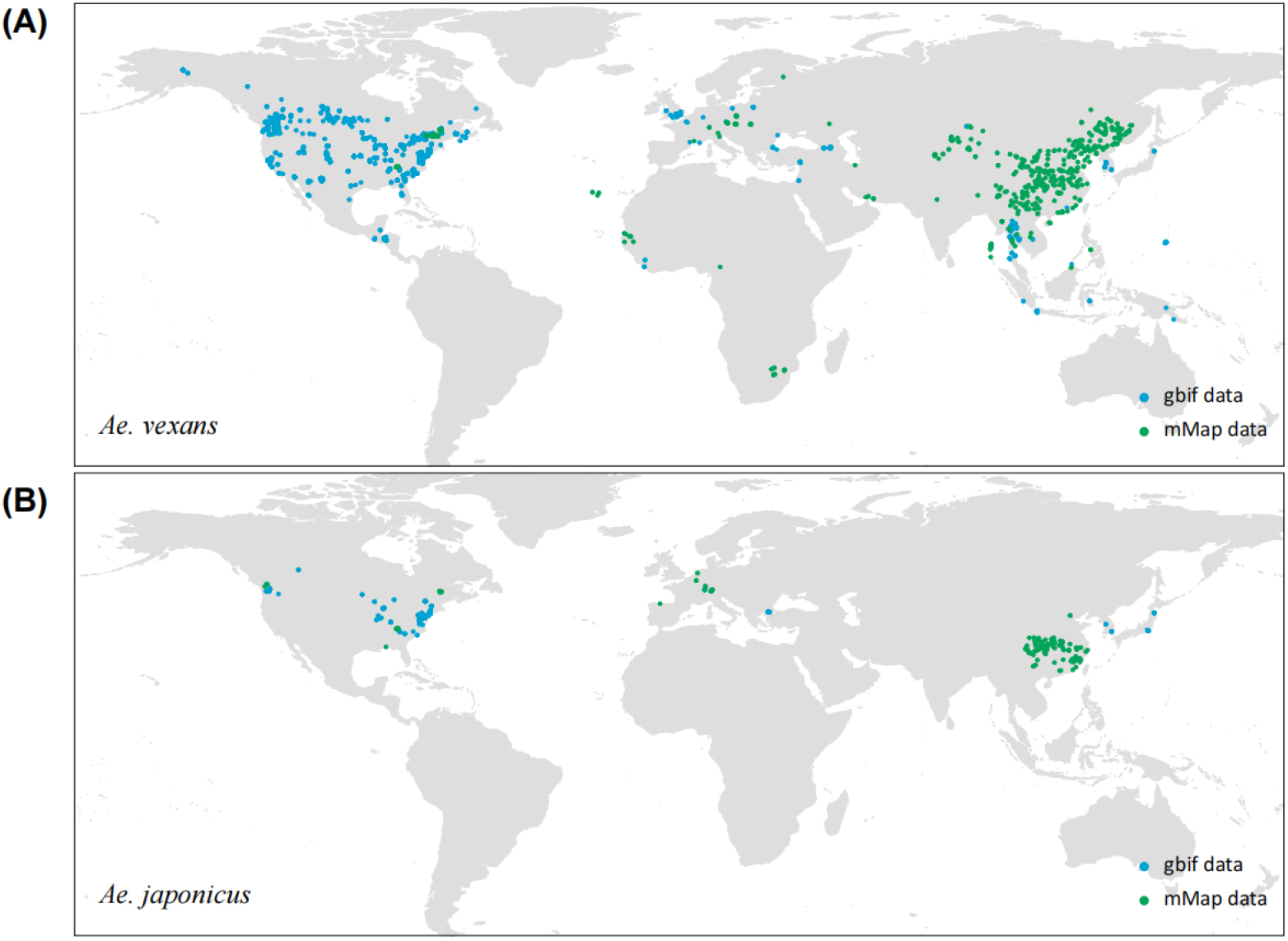
Comparative analysis of mosquito distribution records between GBIF and mMap. (A) Case study of *Aedes vexans*, where mMap data expanded its known distribution by 22 countries/regions (e.g., West Africa, Southern Africa, Northeast Asia, and Europe), significantly revising SDM-predicted ranges. (B) Case study of *Ae. japonicus*.

### 3.3 Several species of the Culicidae family have spread rapidly globally

In recent years, driven by climate change, rapid urbanization, and intensified international trade, several species within the mosquito family (Culicidae) have expanded their global distribution at an unprecedented pace. These mosquitoes thrive in human-altered environments, such as water-holding containers and discarded tires, while rising temperatures and shifting precipitation patterns have facilitated their establishment in temperate regions previously considered unsuitable. A semi-quantitative analysis of occurrence records revealed 28 invasive mosquito species with substantial documentation (Fig. 3). The five most frequently reported species were *Ae. albopictus, Ae. koreicus, Ae. japonicus, A. stephensi*, and *Ae. aegypti* (Fig. 3A). Spatial analysis of invasion sites for these species indicates that Europe has emerged as a major hotspot for mosquito invasions, accounting for a disproportionately high share of documented introductions (Fig. 3B).

**Figure 3.**
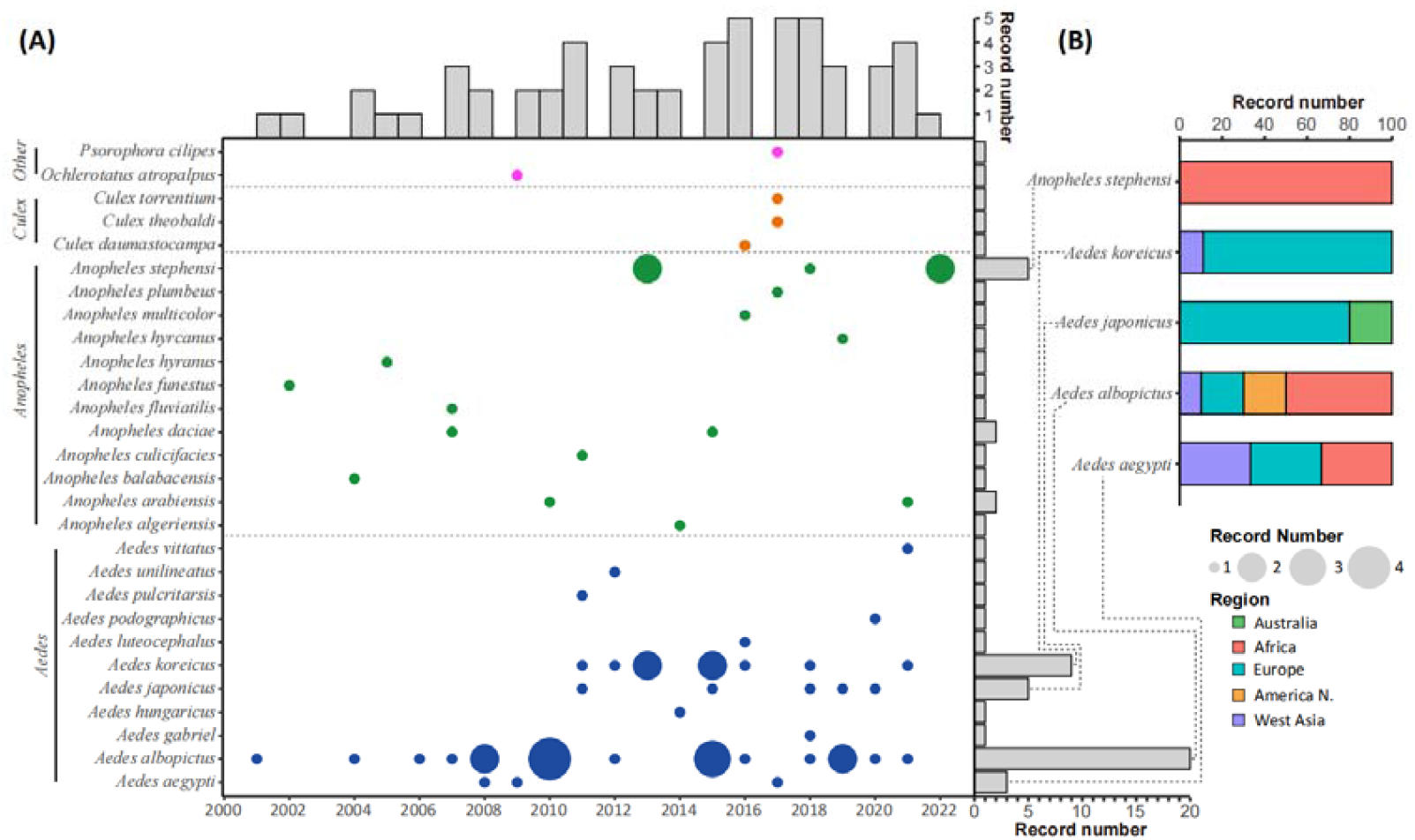
Global patterns of invasive mosquito species. (A) Geographic hotspots of documented invasions, with regions color-coded by the proportion of total introductions. Data derived from semi-quantitative analysis of mMap records. (B) The top five most frequently reported invasive mosquito species, ranked by occurrence records (*Aedes albopictus, Ae. koreicus, Ae. japonicus, Anopheles stephensi*, and *Ae. aegypti*).

### 3.4 Web interface in mMap

mMap provides a user-friendly web interface which enables users to efficiently query, browse, scrutinize, and explore data of mosquito distribution. The interface features an intuitive navigation bar that guides users to key sections, including “Mosquito”, “Database”, “Map”, “Prediction” and “Resources” (Fig. 4A). The “Mosquito” page includes expertly curated knowledge resources on mosquitoes, such as experimental protocols, species identification guidelines, species atlases, and phylogenetic trees. (Fig. 4A). The “Database” page streamlines the process of retrieving data, allowing users to either download the entire dataset or obtain tailored, filtered results with ease (Fig. 4B). The “Map” page further equips users to visualize database entries geographically by applying user-defined filtering criteria (Fig. 4C). The “Resources” section provides important data resource links, including databases on mosquito species knowledge, specialized research platforms, and mosquito distribution information (Fig. 4D). The “Prediction” page features a dedicated web interface for the direct visualization of automated and accelerated species distribution forecasts (Fig. 4A).

**Figure 4.**
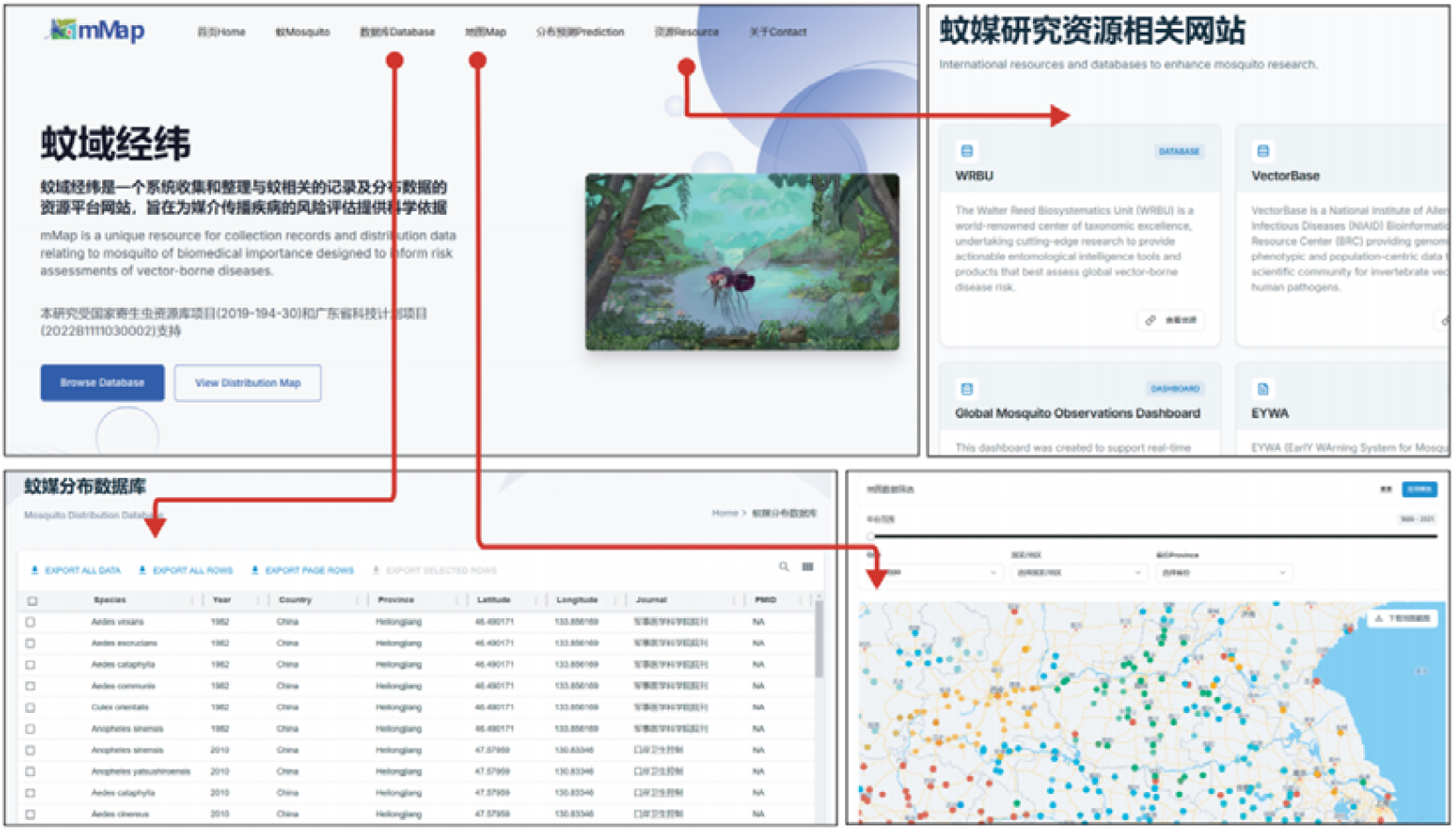
The user interface in mMap website.

## 4. Discussion

The mMap database represents a advancement in consolidating fragmented mosquito distribution data into a unified, accessible resource. By integrating multi-source records from literature, museum collections, and specimen databases across multiple languages and regions, this study addresses critical gaps in understanding mosquito biodiversity and global distribution patterns. The inclusion of previously underrepresented species and regions demonstrates how mMap supplements existing knowledge. For instance, the 22 newly documented countries/regions for *Ae. vexans* highlight how traditional databases like GBIF may underestimate species ranges, leading to biased ecological modeling outcomes.

The rapid global expansion of invasive species such as *Ae. albopictus, Ae. japonicus*, and *A. stephensi* underscores the urgency of monitoring distribution shifts driven by climate change, urbanization, and international trade. Europe’s emergence as a hotspot for mosquito invasions, as evidenced by our semi-quantitative analysis, aligns with recent studies on climate-mediated habitat suitability changes. Conversely, the decline of *Ae. aegypti* in southern China illustrates how local ecological dynamics can counter global trends, necessitating region-specific surveillance strategies. These patterns highlight the dual role of mMap: as a tool for tracking invasions and as a platform to study ecological drivers of distributional changes.

Future research should leverage mMap’s infrastructure to investigate macroecological questions, such as the interplay between mosquito diversity, climate gradients, and disease transmission risks. Expanding the database to include temporal trends and ecological traits (e.g., breeding site preferences, host preferences) could further enhance its utility for mechanistic modeling. Ultimately, mMap’s open-access design and iterative updating mechanism aim to foster global collaboration, bridging gaps between entomology, epidemiology, and conservation biology to mitigate the growing threat of mosquito-borne diseases.

## 5. Conclusion

The mMap database represents a leap in mosquito biodiversity research by integrating multi-source global data to create the most comprehensive distribution resource available, revealing significant range expansions for key vectors that challenge previous estimates from single-source databases. Our findings demonstrate alarming invasion trends, particularly in Europe, driven by climate change and globalization, while highlighting how localized declines like Aedes aegypti in southern China reflect complex ecological dynamics. As an open-access, dynamically updated resource, mMap bridges taxonomic research and public health needs, offering a vital foundation for evidence-based strategies against mosquito-borne diseases in our rapidly changing world.

## CRediT authorship contribution statement

Conceptualization: Xiaohong Zhou and Xiang Guo; Data curation: Xiang Guo; Formal analysis: Xiang Guo, Xiaohua Liu, Liu Ge, Qing He; Funding acquisition: Xiaohong Zhou and Xiang Guo; Investigation: Xiang Guo, Ziyao Li, Shu Zeng, Haiyang Chen, Xiaohua Liu, Liu Ge, Qing He, and Wenyi Li; Methodology: Xiang Guo; Project administration: Xiaohong Zhou; Resources: Xiaohong Zhou; Software: Xiang Guo; Supervision: Xiaohong Zhou; Validation: Ziyao Li, Shu Zeng, Haiyang Chen; Visualization: Xiang Guo; Writing - original draft: Xiang Guo; Writing - review & editing: Xiaohong Zhou.

## Ethics approval and consent to participate

Not applicable.

## Consent for publication

Not applicable.

## Availability of data and materials

The full study protocol and the datasets, are available, following manuscript publication, upon request from the corresponding author.

## Acknowledgements

Not applicable.

## References

1. Soghigian J, Andreadis TG, Livdahl TP. From ground pools to treeholes: convergent evolution of habitat and phenotype in Aedes mosquitoes. BMC Evol Biol. 2017;17 1:262; doi: 10.1186/s12862-017-1092-y.

2. Reidenbach KR, Cook S, Bertone MA, Harbach RE, Wiegmann BM, Besansky NJ. Phylogenetic analysis and temporal diversification of mosquitoes (Diptera: Culicidae) based on nuclear genes and morphology. BMC Evol Biol. 2009;9:298; doi: 10.1186/1471-2148-9-298.

3. da Silva AF, Machado LC, de Paula MB, da Silva Pessoa Vieira CJ, de Morais Bronzoni RV, de Melo Santos MAV, et al. Culicidae evolutionary history focusing on the Culicinae subfamily based on mitochondrial phylogenomics. Sci Rep. 2020;10 1:18823; doi: 10.1038/s41598-020-74883-3.

4. Guzman MG, Gubler DJ, Izquierdo A, Martinez E, Halstead SB. Dengue infection. Nat Rev Dis Primers. 2016;2:16055; doi: 10.1038/nrdp.2016.55.

5. Petersen LR, Jamieson DJ, Powers AM, Honein MA. Zika Virus. N Engl J Med. 2016;374 16:1552–63; doi: 10.1056/NEJMra1602113.

6. Franklinos LHV, Jones KE, Redding DW, Abubakar I. The effect of global change on mosquito-borne disease. Lancet Infect Dis. 2019;19 9:e302–e12; doi: 10.1016/S1473-3099(19)30161-6.

7. Li Y, Kamara F, Zhou G, Puthiyakunnon S, Li C, Liu Y, et al. Urbanization increases Aedes albopictus larval habitats and accelerates mosquito development and survivorship. PLoS Negl Trop Dis. 2014;8 11:e3301; doi: 10.1371/journal.pntd.0003301.

8. van Klink R, Bowler DE, Gongalsky KB, Swengel AB, Gentile A, Chase JM. Meta-analysis reveals declines in terrestrial but increases in freshwater insect abundances. Science. 2020;368 6489:417–20; doi: 10.1126/science.aax9931.

9. Kaufman MG, Fonseca DM. Invasion biology of Aedes japonicus japonicus (Diptera: Culicidae). Annu Rev Entomol. 2014;59:31–49; doi: 10.1146/annurev-ento-011613-162012.

10. Bonizzoni M, Gasperi G, Chen X, James AA. The invasive mosquito species Aedes albopictus: current knowledge and future perspectives. Trends Parasitol. 2013;29 9:460–8; doi: 10.1016/j.pt.2013.07.003.

11. Kraemer MUG, Reiner RC, Jr., Brady OJ, Messina JP, Gilbert M, Pigott DM, et al. Past and future spread of the arbovirus vectors Aedes aegypti and Aedes albopictus. Nat Microbiol. 2019;4 5:854–63; doi: 10.1038/s41564-019-0376-y.

12. Zhao M, Ran X, Zhang Q, Gao J, Wu M, Xing D, et al. Genetic diversity of Flaviviridae and Rhabdoviridae EVEs in Aedes aegypti and Aedes albopictus on Hainan Island and the Leizhou Peninsula, China. Infect Genet Evol. 2024;123:105627; doi: 10.1016/j.meegid.2024.105627.

13. Liu P, Lu L, Jiang J, Guo Y, Yang M, Liu Q. The expanding pattern of Aedes aegypti in southern Yunnan, China: insights from microsatellite and mitochondrial DNA markers. Parasit Vectors. 2019;12 1:561; doi: 10.1186/s13071-019-3818-8.

14. Lv RC, Zhu C, Wang CH, Ai LL, Lv H, Zhang B, et al. Genetic diversity and population structure of Aedes aegypti after massive vector control for dengue fever prevention in Yunnan border areas. Sci Rep. 2020;10 1:12731; doi: 10.1038/s41598-020-69668-7.

15. Kass JM, Fukaya K, Thuiller W, Mori AS. Biodiversity modeling advances will improve predictions of nature’s contributions to people. Trends Ecol Evol. 2024;39 4:338–48; doi: 10.1016/j.tree.2023.10.011.

16. Kass JM, Guenard B, Dudley KL, Jenkins CN, Azuma F, Fisher BL, et al. The global distribution of known and undiscovered ant biodiversity. Sci Adv. 2022;8 31:eabp9908; doi: 10.1126/sciadv.abp9908.

